# Mitochondrial respiration modulates Hsf1 activation and the heat shock response

**DOI:** 10.64898/2026.05.07.723568

**Authors:** D.W. McDonald, A. Dea, R. Sava, Y.J. Kim, L. Joos, D. Pincus, M.L Duennwald

## Abstract

Cells employ a bevy of transcriptional and post-translational stress responses to tolerate the burden of misfolded proteins induced by stress. In particular, the heat shock response facilitates the upregulation of molecular chaperones and protein remodeling factors that mediate proteostasis in response to accumulated misfolded proteins in the nucleus and cytosol. However, in response to stress neurons struggle to induce a canonical heat shock response, highlighting our poor understanding of how neurons maintain proteostasis. Specifically, the ability of post-mitotic respiring cells to regulate the heat shock response in comparison to their rapidly dividing, predominantly glycolytic counterparts has been under-studied. In this study, we employ yeast models that are easily manipulated to generate energy via glycolysis or mitochondrial respiration by changing the carbon source in the media. Using this model, we demonstrate that Hsf1 activity, the heat shock response and proteostasis are impaired in respiring cells. Interestingly, our data show that reduced Hsf1 activity regulates viability of respiring cells, with respiring cells poorly tolerating constitutively activated Hsf1. Finally, we describe alternative post-translational programming of the molecular chaperones Hsp70 and Hsp104 that plausibly enables respiring cells to mediate proteostasis despite a dampened heat shock response. Our findings offer new insights into possible proteostatic strategies employed by cells in different metabolic conditions.

## INTRODUCTION

Cells respond to the accumulation of misfolded proteins through a set of transcriptional programs called stress responses that regulate the expression of molecular chaperones and protein remodeling factors to mediate proteostasis [1–4]. Cytosolic and nuclear proteostasis is predominantly mediated by the heat shock response [1]. Misfolding of cytosolic and nuclear proteins activates Heat shock factor 1 (Hsf1), which binds to heat shock elements in the promotor regions of genes encoding heat shock proteins and molecular chaperones to upregulate their transcription [5–10]. These molecular chaperones restore proteostasis by refolding misfolded proteins or targeting them for degradation [4,11–18]. The heat shock response has been extensively studied in glycolytic cells, and has been shown to regulate development, proliferation and stress tolerance, including regulating protein aggregation associated with ageing and neurodegenerative diseases [7,8,10,14,17,19–28]. Yet, targeting molecular chaperones for the maintenance of proteostasis in neurodegenerative diseases has not proved fruitful thus far, highlighting our particularly poor understanding of how proteostasis is regulated in neurons.

Adenosine triphosphate (ATP) is a crucial energy source used to fuel essential cellular functions. Cellular ATP is generated through various metabolic pathways, chiefly by converting glucose to pyruvate via glycolysis, or metabolizing acetyl-coA to liberate electrons that are shuttled through the electron transport chain to generate ATP via oxidative phosphorylation [29–31]. Glycolytic strategies to generate ATP are often adopted by rapidly dividing cells, as glycolysis generates ATP rapidly and yields many macromolecules essential for proliferation [30,32,33]. In contrast, cells with high energy demands (i.e., neurons) preferentially generate energy via the more efficient oxidative phosphorylation [34,35]. Notably, each metabolic program poses unique challenges for cells, with glycolysis increasing extracellular pH through the production of lactic acid and oxidative phosphorylation inducing oxidative stress via the production of free radicals [36–39]. In addition, metabolic program may modify the pool of ATP available in cells, challenging the maintenance of folded proteins facilitated either by direct interactions with ATP or by ATP-dependent molecular chaperones [40–43].

Yeast cells provide an ideal model organism to investigate how metabolism regulates the heat shock response and cytosolic proteostasis. In typical laboratory conditions, yeast cells ferment glucose via glycolysis to generate ATP [44,45]. However, when grown in the absence of fermentable carbon, yeast cells adapt to generate energy via oxidative phosphorylation [46]. Metabolic manipulation of yeast has provided key insights into how mitochondrial function regulates ageing and tolerance of neurodegenerative disease-associated misfolded proteins [47–52]. Additionally, the heat shock response and its role in cytosolic protein quality control has been extensively described in yeast [5–10,12,15,16,23,25–28,53].

Here, we take advantage of the ease with which metabolism can be manipulated in yeast cells to determine how metabolic activity changes cellular responses to stress. Specifically, we hypothesize that different metabolic programs modulate the heat shock response and, in turn, proteostasis. Our data reveals that Hsf1 activation is impaired in respiring cells, leading to a dampened heat shock response and a reduced capacity to maintain proteostasis. Additionally, we demonstrate that the activity of Hsf1 regulates the viability of respiring cells, whereby increased Hsf1 activity is poorly tolerated by respiring cells. Furthermore, we describe functional reprogramming of the molecular chaperones Hsp104 and Hsp70 in respiring cells, which enable cells to adapt to the proteostatic demands induced by respiration. Taken together, our findings highlight a need to interrogate how energy metabolism regulates proteostasis. Notably, our findings demonstrate a plausible mechanism by which neurons may be more susceptible to protein misfolding and impaired proteostasis in age-related neurodegenerative diseases due to their reliance on mitochondrial respiration for ATP production.

## RESULTS

### The heat shock response is severely dampened in respiring cells

The heat shock response is a pro-survival transcriptional program elicited by protein misfolding in the nucleus and/or cytosol [1,10,14]. Although the heat shock response is well described in glycolytic, rapidly dividing cells, its regulation in senescent cells that favor oxidative phosphorylation, such as neurons, is unclear. To better understand how metabolic program regulates the heat shock response, we took advantage of the ease at which metabolic activity is manipulated in yeast through exposure to different carbon sources. Under standard laboratory conditions, yeast cells ferment glucose to generate ATP via glycolysis [44,45]. However, when forced to metabolize non-fermentable carbon (i.e., glycerol), yeast cells undergo a series of transcriptional changes that facilitate the switch towards mitochondrial respiration for energy production [46]. We used this paradigm to compare the transcriptional profile of glycolytic and respiring cells exposed to a 30-minute heat shock at 42°C.

Our principal component analysis indicates distinct transcriptional profiles across treatments, with principal component 1 reflecting variability associated with temperature upshift, and principal component 2 reflecting variability associated with metabolic program (Fig. 1A). Heat shock induces mostly upregulated genes in both glycolytic and respiring cells, although the number of upregulated genes and the amplitude of their upregulation is higher in glycolytic cells (Fig. 1B-C). The genes upregulated (log2FC>3) by heat shock in both glycolytic and respiring cells are enriched for gene ontology (GO) terms related to proteostasis and are regulated by Hsf1, the transcriptional regulator of the heat shock response [5,6,9] (Fig. 1D-E). However, the enrichment of proteostasis-related and Hsf1-regulated genes within the set of genes upregulated by heat shock is most pronounced in glycolytic cells (Fig. 1D-E). Additionally, heat shock upregulates genes associated with cell morphogenesis in respiring cells, but not in glycolytic cells (Fig. 1D-E). When examining the normalized counts for candidate genes known to be upregulated by the heat shock response (HSP104, HSP42, HSP82, and SSA1), we find that these genes are upregulated by heat shock to a greater extent in glycolytic cells when compared to respiring cells (Fig. 1F-I). Interestingly, this upregulation was not observed for all heat shock-regulated genes, as SSA3 is upregulated to the same degree by heat shock in respiring cells when compared to glycolytic cells (Fig. 1J).

**Figure 1:**
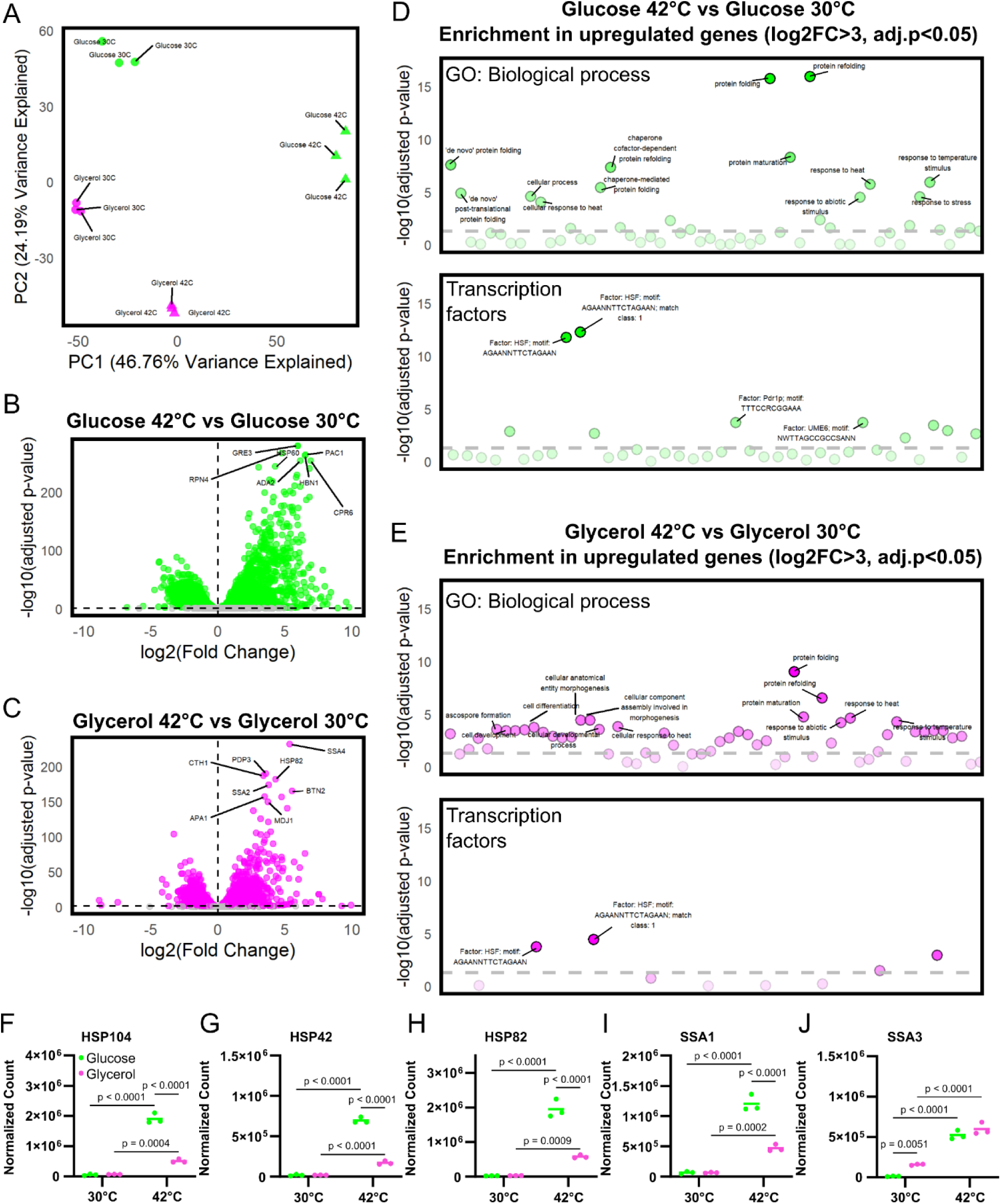
Mitochondrial respiration dysregulates transcriptional response to heat shock. A. PCA plot of transcriptional profile of cells grown in glucose or glycerol and either grown at 30°C or heat shocked at 42°C for 30 minutes. B. Volcano plot showing upregulated and downregulated genes in cells grown in glucose and heat shocked at 42°C for 30 mins relative to cells grown in glucose at 30°C. C. Volcano plot showing upregulated and downregulated genes in cells grown in glycerol and heat shocked at 42°C for 30 mins relative to cells grown in glycerol at 30°C. D. Enriched GO terms (Biological processes) and transcription factors associated with the set of upregulated genes (log2FC>3) in cells grown in glucose and heat shocked at 42°C for 30 mins relative to cells grown in glucose at 30°C. E. Enriched GO terms (Biological processes) and transcription factors associated with the set of upregulated genes (log2FC>3) in cells grown in glycerol and heat shocked at 42°C for 30 mins relative to cells grown in glycerol at 30°C. F-J. Normalized transcript counts for heat shock genes F. HSP104, G. HSP42, H. HSP82, I. SSA1 and J. SSA3 in cells grown in glucose (green) or glycerol (magenta) and either grown at 30°C or heat shocked at 42°C for 30 minutes.

We next compared transcriptional profiles of respiring and glycolytic cells at the same temperature (i.e., 30°C or 1-hour heat shock at 42°C). In cells grown at 30°C, glycerol metabolism induces a similar distribution of upregulated and downregulated genes (Fig. S1A). The genes upregulated (log2FC>2.2) by glycerol metabolism in cells grown at 30°C are enriched for GO terms associated with fatty acid metabolism and are regulated by Ume6, a transcriptional repressor that regulates oxidative metabolism and meiosis [54,55], and Cat8, a transcriptional activator that regulates oxidative metabolism [56,57] (Fig. S1B). The genes downregulated (log2FC<-2.2) by glycerol metabolism in cells grown at 30°C are enriched for GO terms associated with transmembrane transport and are regulated by Xbp1, a transcriptional activator that regulates quiescence [58,59], and Rgt1, a transcription factor that regulates expression of glucose transporter genes [60,61] (Fig. S1.1C). In heat shocked cells, glycerol metabolism induces mostly downregulated genes (Fig. S1D). Interestingly, the genes upregulated (log2FC>2.2) by glycerol metabolism in heat shocked cells are enriched for GO terms associated with ribosome biogenesis (Fig. S1E). The genes downregulated (log2FC<-2.2) by glycerol metabolism in heat shocked cells are enriched for GO terms associated with transmembrane transport and are regulated by the transcription factors Xbp1, Rgt1, and Rsc3/30, components of the RSC chromatin remodeling complex, which regulates expression of ribosomal protein genes [62,63] (Fig. S1F).

Because heat shocked respiring cells show upregulation in ribosome biogenesis related genes when compared to heat shocked glycolytic cells, we opted to monitor protein biosynthesis. Using a click-chemistry based approach [64,65], we pulse labelled de novo synthesized proteins in glycolytic and respiring cells before and after an hour heat shock at 42°C. We confirm previous findings showing that heat shock inhibits protein biosynthesis in glycolytic cells [66]. By contrast, heat shock has no effect on protein biosynthesis in respiring cells (Fig. S1G-I).

In sum, respiring cells mount an overall weaker transcriptional response to heat shock when compared to glycolytic cells. In addition, respiring cells show increased levels of transcripts associated with ribosome biogenesis and increased de novo synthesized protein levels after heat shock when compared to glycolytic cells.

### Impaired activation of the heat shock response condensate cascade in respiring cells

At the cell biological level, the heat shock response is mediated via a condensate cascade in which nucleolar stress arrests ribosome biogenesis and results in the accumulation of orphan ribosomal proteins (oRPs), which condense on the surface of the nucleolus and recruit the chaperones Sis1 and Hsp70 away from the nucleoplasm where they repress Hsf1 [67,68]. Hsf1 is then released from Hsp70 and itself condenses with the transcriptional machinery to drive HSR gene expression [69,70]. To determine whether the carbon source influences this signaling cascade, we imaged the components in budding cells grown in glucose (YPD) or glycerol (YP-glycerol) medium with and without heat shock (39°C, 15 min). In glucose grown cells, Hsf1-mKate formed condensates upon heat shock, as reflected by an increase in the Hsf1 condensate index (Figure 2A, 2C; p < 0.01). In glycerol-grown cells, Hsf1 condensate formation was attenuated, and the Hsf1 signal overlapped with mitochondrial autofluorescence, suggesting altered localization that may reflect a non-nuclear pool of Hsf1 (Figure 2A).

**Figure 2:**
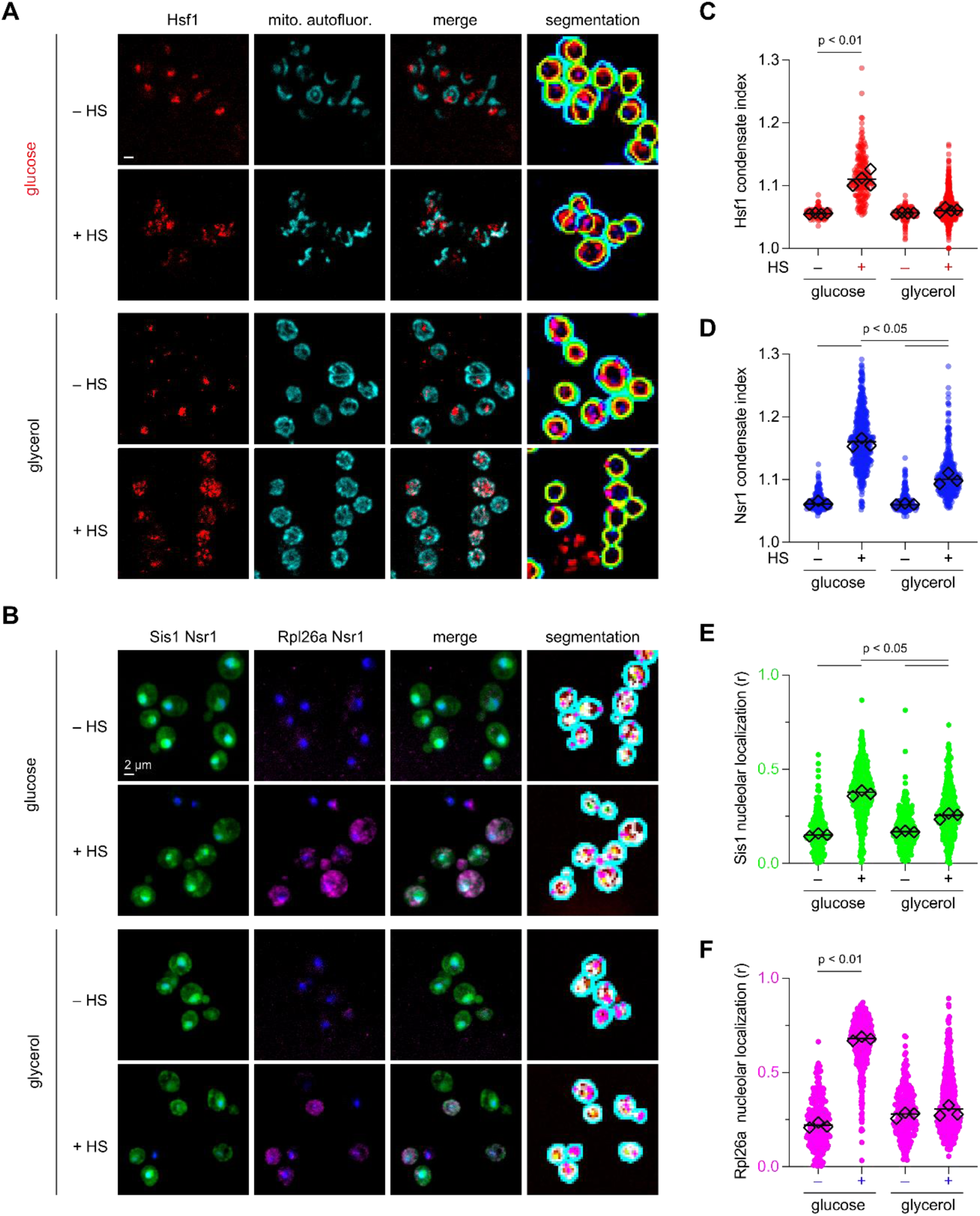
Carbon source modulates Hsf1 condensate formation and nucleolar organization during heat shock. A. Representative fluorescence images of budding yeast expressing Hsf1-mKate (red) grown in glucose (top) or glycerol (bottom) medium, with or without heat shock (39°C, 15 min). Mitochondrial autofluorescence is shown in cyan. Left to right: Hsf1, mitochondrial autofluorescence, merge, cellquant segmentation overlay with cell boundaries. Scale bar, 2 µm. Strain: DPY430 (W303 MATa HSF1-mKate::URA3). B. Representative fluorescence images of a separate strain expressing Sis1-mVenus (green), Rpl26a-HaloTag pulse-labeled with JF-646 (magenta), and Nsr1-mScarlet-I (blue, nucleolar marker), under the same conditions as A. Left to right: Sis1/Nsr1 overlay, Rpl26a/Nsr1 overlay, merge, cellquant segmentation overlay. Scale bar, 2 µm. Strain: DPY1766 (W303 MATa SIS1-mVenus::HIS3 RPL26A-HALO NSR1-mScarlet-I). C. Hsf1 condensate index (95th percentile / mean pixel intensity) across conditions. Each small dot represents a single cell; large diamonds indicate replicate medians (n ≥ 3 biological replicates per condition, >200 cells per replicate). P-values from Wilcoxon rank-sum test on replicate medians. D. Nsr1 condensate index (95th percentile / mean pixel intensity) across conditions (p < 0.05). Plotting conventions as in C. E. Sis1–Nsr1 Pearson colocalization coefficient (r), quantifying Sis1 nucleolar localization. Plotting conventions as in C. F. Rpl26a–Nsr1 Pearson colocalization coefficient (r), quantifying nucleolar localization of newly synthesized (pulse-labeled) Rpl26a. Plotting conventions as in C.

Upstream of Hsf1 condensation, heat shock drives the accumulation of oRPs around the nucleolus, which recruits the chaperone Sis1 away from Hsf1 [67]. To determine whether carbon source affects this pathway, we imaged Sis1-mVenus, Rpl26a-HaloTag, and Nsr1-mScarlet-I in a single strain co-expressed from their endogenous loci. We pulse labeled with JF-646 to label newly synthesized Rpl26a as a proxy for orphan ribosomal proteins (oRPs) in cells grown in glucose and glycerol in the absence and presence of heat shock. The nucleolar marker Nsr1 exhibited carbon source dependent changes in its condensate index upon heat shock, with reduced condensation in glycerol (Figure 2D; p < 0.05). Sis1 nucleolar colocalization increased upon heat shock in glucose grown cells to a greater extent than in glycerol (Figure 2E; p < 0.05). Newly synthesized pulse-labeled Rpl26a also showed high nucleolar retention upon heat shock in glucose, which was nearly obviated in glycerol (Figure 2F). Together, these results reveal that carbon source modulates the nucleolar response to heat shock, affecting nucleolar morphology, the localization dynamics of oRPs and Sis1, and Hsf1 condensate formation.

### Respiring cells are deficient at restoring proteostasis

Because we show that respiring cells exhibit a deficient heat shock response, we tested whether this influenced the ability of respiring cells to maintain proteostasis. To this end, we expressed a metastable protein, firefly luciferase-GFP (FFL-GFP), which readily forms aggregates upon exposure to stress [52], in cells as a reporter to monitor the state of proteostasis in cells generating energy by different metabolic programs.

We first performed growth assays at 30°C to assess whether FFL-GFP toxicity is exacerbated by metabolism of non-fermentable carbon sources. FFL-GFP induces a growth defect in cells generating energy by glycolysis (glucose) (Fig. 3A). This growth defect is unchanged in cells generating energy by oxidative phosphorylation (glycerol and KoAc) or by β-oxidation (myristic acid and oleic acid) (Fig. 3A). We next tested whether FFL-GFP aggregation is induced by metabolism of non-fermentable carbon sources. FFL-GFP does not form fluorescent foci in cells metabolizing glucose, glycerol, KoAc, myristic acid or oleic acid (Fig. 3B).

**Figure 3:**
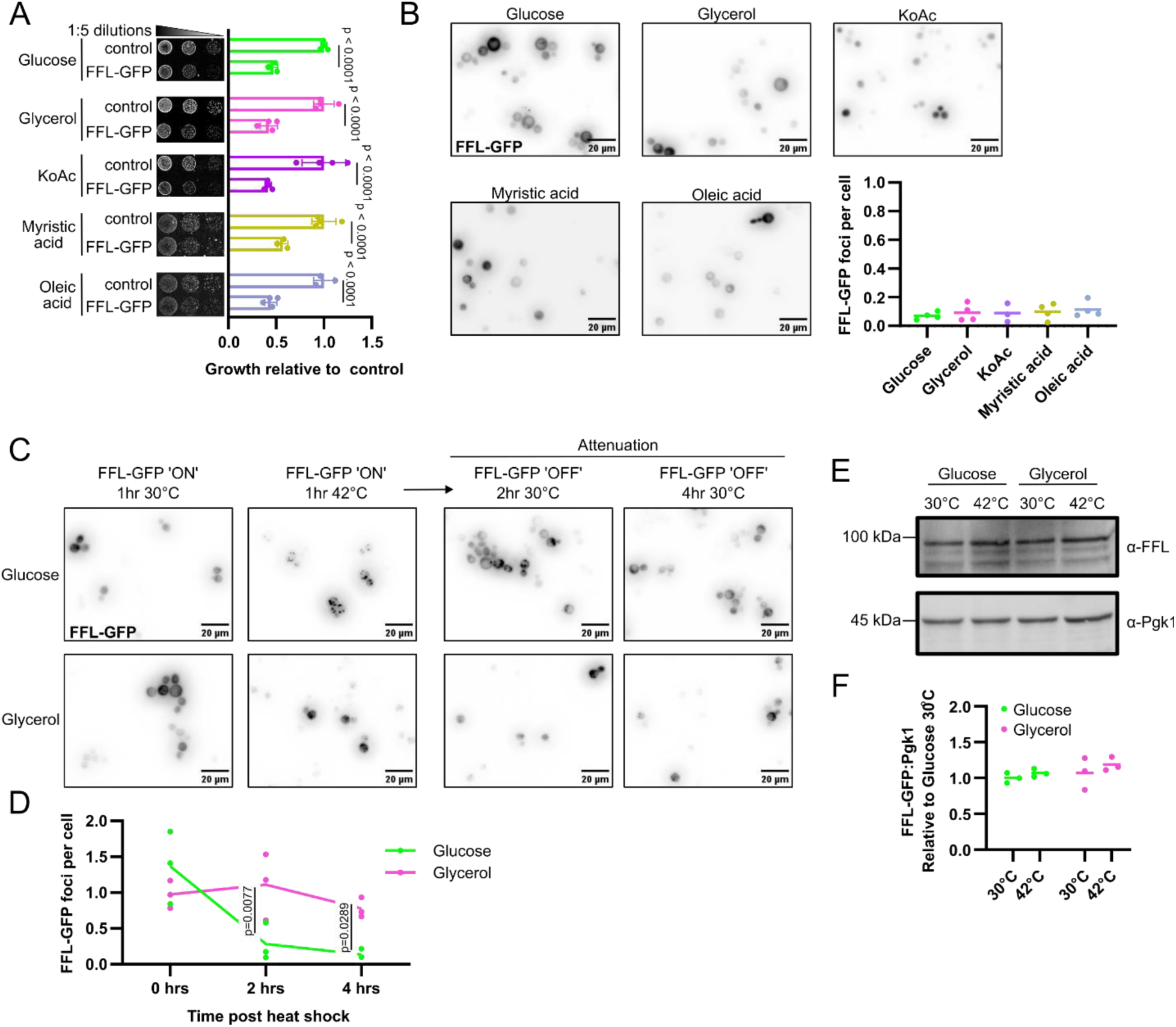
Impaired aggregate clearance in respiring cells. A. Growth assays of cells expressing either vector control or FFL-GFP on plates supplemented with either glucose, glycerol, potassium acetate (KoAc), myristic acid or oleic acid at 30°C. B. Fluorescence microscopy of cells expressing FFL-GFP grown in media supplemented with either glucose, glycerol, potassium acetate (KoAc), myristic acid or oleic acid. C. Fluorescence microscopy and D. quantification showing the persistence of FFL-GFP fluorescent foci in cells grown in media supplemented with glucose or glycerol following a one-hour heat shock at 42°C and recovery at 30°C for two or four hours. E. Western blot and F. quantification showing the steady state levels of FFL-GFP in cells grown and heat shocked in media supplemented with glucose or glycerol.

Although mitochondrial respiration does not induce aggregation of FFL-GFP on its own, we speculated that an impaired heat shock response in respiring cells would inhibit the clearance of aggregated FFL-GFP. To test this hypothesis, we induced the formation of FFL-GFP aggregates in glycolytic and respiring cells via a 1-hour heat shock at 42°C. We show that a heat shock does not change the number of FFL-GFP aggregates in respiring cells when compared to glycolytic cells (Fig. 3C-D). After inducing aggregation of FFL-GFP, we monitored the dissolution of FFL-GFP aggregates while the cells recover at 30°C. We find that glycolytic cells rapidly dissolve FFL-GFP aggregates after 2 hours (Fig. 3C-D). However, respiring cells are delayed in their dissolution of FFL-GFP aggregates (Fig. 3C-D). Furthermore, we show that the changes in FFL-GFP aggregate dispersal are not associated with a change in steady-state levels of FFL-GFP, as FFL-GFP is expressed at equal levels in glycolytic and respiring cells (Fig. 3E-F).

In sum, fermenting cells rapidly resolve stress-induced aggregation of FFL-GFP, whereas respiring cells are less effective at doing so.

### Viability and proteostasis in respiring cells depend on HSF1 activity

Because we found a correlation between a dampened heat shock response and the inability to restore proteostasis following heat shock, we next assessed whether Hsf1 activity regulates viability and proteostasis in respiring cells. To this end, we employed yeast strains deleted for HSF1 at its genomic locus and complemented with either the wild-type HSF1 allele, the overactive *hsf1ΔN* allele or the inhibited *hsf1ΔC* allele [8].

We first performed growth assays to determine whether genetic manipulation of Hsf1 activity affects the growth behavior of respiring cells. Our data demonstrates that *hsf1ΔN* is well tolerated while *hsf1ΔC* induces a strong growth defect in glycolytic cells (glucose) grown at 30°C (Fig. 4A). Interestingly, the inverse is true in respiring cells (glycerol) grown at 30°C, with *hsf1ΔN* inducing a strong growth defect and *hsf1ΔC* being well tolerated (Fig. 4A). The growth defects induced by *hsf1ΔC* in glycolytic cells and *hsf1ΔN* in respiring cells are exacerbated by elevated temperature (Fig. 4B). Notably, the expression of FFL-mCherry had no effect on growth, regardless of the strain or growth conditions (Fig 4A-B).

**Figure 4:**
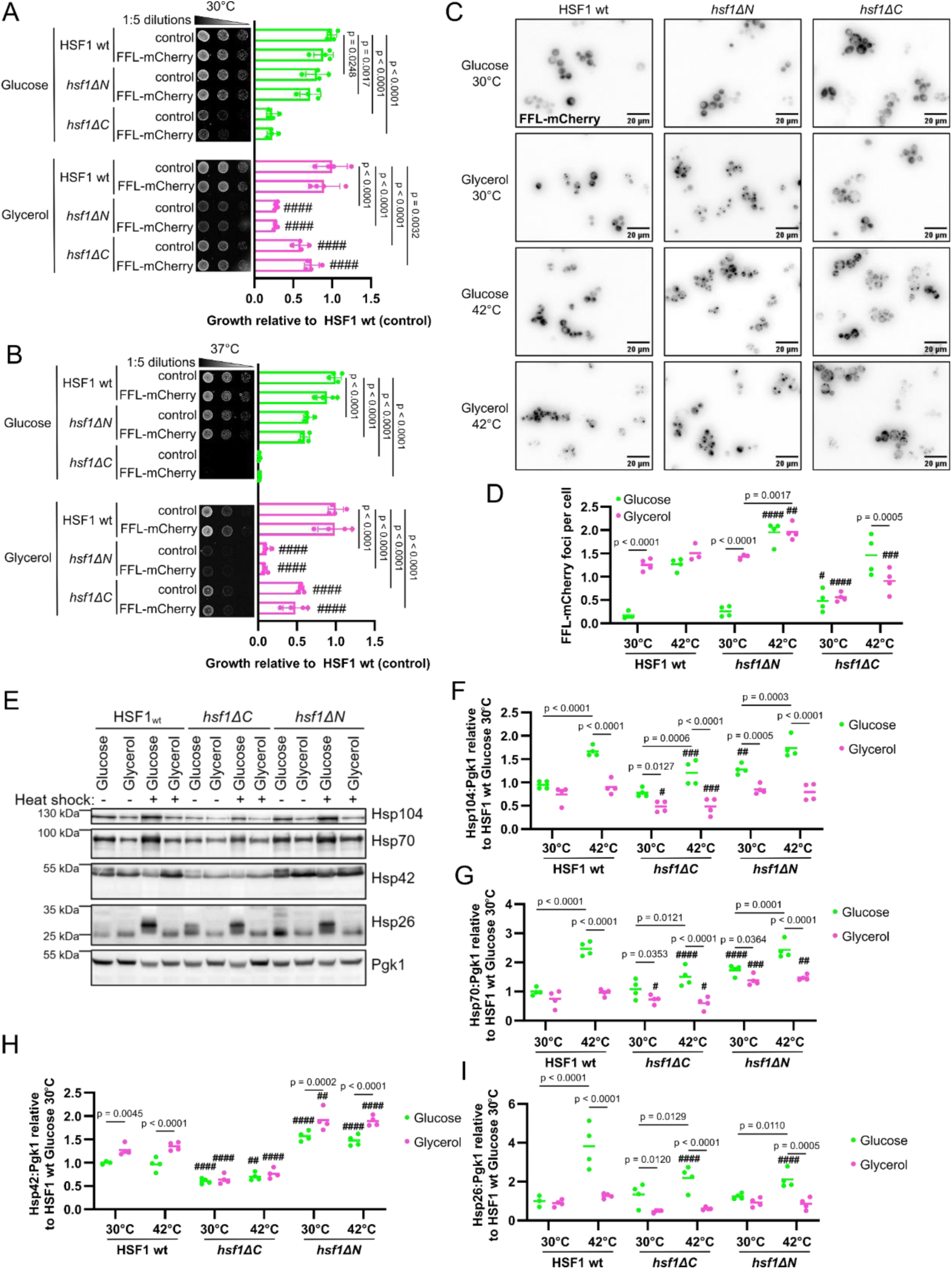
HSF1 activity regulates viability of respiring cells. Growth assays of HSF1 wt, *hsf1ΔN*, and *hsf1ΔC* cells expressing either vector control or FFL-mCherry on plates supplemented with either glucose or glycerol and incubated at A. 30°C or B. 37°C. The mean growth (+/- SD) relative to HSF1 wt control cells are represented graphically for at least three independent experiments. # symbols indicate significant differences compared to the same sample metabolizing glucose at the same temperature (*# p<0.05, ## p<0.01, ### p<0.001, #### p<0.0001*). C. Fluorescence microscopy and D. quantification showing the number of FFL-mCherry foci per cell of HSF1 wt, *hsf1ΔN*, and *hsf1ΔC* cells grown in media supplemented with either glucose or glycerol at 30°C or heat shocked at 42°C for 1 hour. The number of FFL-mCherry foci per cell is represented graphically for at least three independent experiments. # symbols indicate significant differences compared to HSF1 wt cells metabolizing glucose at the same temperature (*# p<0.05, ## p<0.01, ### p<0.001, #### p<0.0001*).

We next assessed whether HSF1 activity affects proteostasis in respiring cells. To this end, we monitored aggregation of the proteostasis reporter FFL-mCherry. FFL-mCherry is soluble in both HSF1 wt and *hsf1ΔN* cells metabolizing glucose at 30°C, while *hsf1ΔC* induces a moderate increase in FFL-mCherry aggregation in cells metabolizing glucose at 30°C when compared to HSF1 wt cells (Fig. 4C-D). Metabolism of glycerol at 30°C increases aggregation of FFL-mCherry in HSF1 wt and *hsf1ΔN* cells when compared to cells metabolizing glucose at 30°C (Fig. 4C-D). By contrast, *hsf1ΔC* cells grown in glycerol at 30°C show no change in FFL-mCherry aggregation compared to *hsf1ΔC* cells grown in glucose (Fig. 4C-D). Furthermore, *hsf1ΔC* cells metabolizing glycerol at 30°C show a marked reduction in FFL-mCherry aggregates when compared to HSF1 wt cells (Fig. 4C-D). A one-hour heat shock induces an increase in FFL-mCherry aggregation in HSF1 wt, *hsf1ΔN* and *hsf1ΔC* cells metabolizing glucose (Fig. 4C-D). However, *hsf1ΔN* induces more FFL-mCherry aggregates in heat shocked cells metabolizing glucose when compared to HSF1 wt cells (Fig. 4C-D). In contrast, a heat shock in cells metabolizing glycerol only increased FFL-mCherry aggregation in *hsf1ΔN* cells, but not HSF1 wt and *hsf1ΔC* cells (Fig. 4C-D). Notably, *hsf1ΔC* heat shocked cells metabolizing glycerol had fewer FFL-mCherry aggregates than HSF1 wt cells (Fig. 4C-D).

Cells mediate proteostasis in response to heat shock through Hsf1-dependent upregulation of many molecular chaperones and protein remodeling factors that mediate proteostasis by dispersing aggregates and refolding misfolded proteins [4,12–17,26–28,71–73]. We sought to determine how the Hsf1 variants interfere with proteoastasis by modulating the expression of molecular chaperones in respiring cells. We performed western blots using lysates extracted from glycolytic and respiring cells subjected to a 42°C heat shock for one hour (Fig. 4E). We demonstrate that Hsp104, Hsp70 and Hsp26, but not Hsp42, increase in steady-state levels in glycolytic HSF1 wt cells after heat shock (Fig. 4F-I) Similar to our transcriptional profiling, we find that Hsp104, Hsp70, Hsp42 and Hsp26 protein levels do not increase in respiring HSF1 wt cells after heat shock (Fig. 4F-I). Notably, Hsp42 protein levels increase in respiring HSF1 wt cells both before and after heat shock when compared to glycolytic HSF1 wt cells. In contrast to glycolytic HSF1 wt cells, Hsp104, Hsp70, Hsp42 and Hsp26 protein levels are reduced in glycolytic *hsf1ΔC* cells after heat shock (Fig. 4F-I). In addition, respiring *hsf1ΔC* cells show reduced Hsp104, Hsp70, Hsp42 and Hsp26 protein levels when compared to respiring HSF1 wt cells both before and after heat shock (Fig. 4F-I). Hsp104, Hsp42 and Hsp70, but not Hsp26, protein steady-state levels are increased in glycolytic *hsf1ΔN* cells before heat shock when compared to glycolytic HSF1 wt cells before heat shock (Fig. 4F-G). However, glycolytic *hsf1ΔN* cells show no increase in Hsp104 and Hsp70 protein levels while Hsp26 protein levels are decreased post heat shock when compared to glycolytic HSF1 wt cells post heat shock (Fig. 4F, 4G & 4I). Generally, *hsf1ΔN* was sufficient to increase Hsp70 and Hsp42, but not Hsp104 and Hsp26, protein levels in respiring cells when compared to HSF1 wt (Fig. 4G-H).

In sum, a decline in Hsf1 activity impairs growth and proteostasis in glycolytic cells but is well tolerated and preserved proteostasis in respiring cells. Conversely, increased Hsf1 activity is well tolerated in glycolytic cells but impairs growth in respiring cells. Interestingly, increased Hsf1 activity increased aggregation of a metastable protein, regardless of metabolic program. Finally, deficient Hsf1 activity reduced the protein levels of Hsp104, Hsp70 and Hsp42 in respiring cells, while increased Hsf1 activity increased the protein levels of Hsp70 and Hsp42.

### Modified function of Hsp104 and Hsp70 in respiring cells

Our previous data strongly indicate that manipulating the expression of molecular chaperones has drastic effects on viability and proteostasis of respiring cells. Thus, we sought to determine how specific molecular chaperones mediate proteostasis in respiring cells. Two major molecular chaperones that mediate proteostasis in response to stress are Hsp104 and Hsp70. Hsp104 is a AAA-ATPase which cooperates with Hsp70 to mediate the dissolution of aggregates [12,16,71,74–76]. In addition, Hsp70 broadly regulates protein refolding in the cytosol and the nucleus [8,11,17,18]. We hypothesized that Hsp104 and Hsp70 regulate the dispersal of protein aggregates in respiring cells, and as such, mediate viability of respiring cells in response to stress.

To test this hypothesis, we investigated whether the functions of Hsp104 and Ssa1 change during stress in respiring cells. To this end, we monitored inclusion formation of Hsp104-YFP and Ssa1-GFP. We find that Hsp104-YFP forms inclusions following a heat shock in both glycolytic and respiring cells (Fig. 5A-B). However, the number of Hsp104-YFP inclusions formed in response to heat shock is increased in respiring cells when compared to glycolytic cells (Fig. 5A-B). Similarly, Ssa1-GFP forms inclusions following a heat shock of glycolytic cells (Fig. 5C-D). However, the inclusion formation of Ssa1-GFP following a heat shock is abolished in respiring cells (Fig. 5C-D).

**Figure 5:**
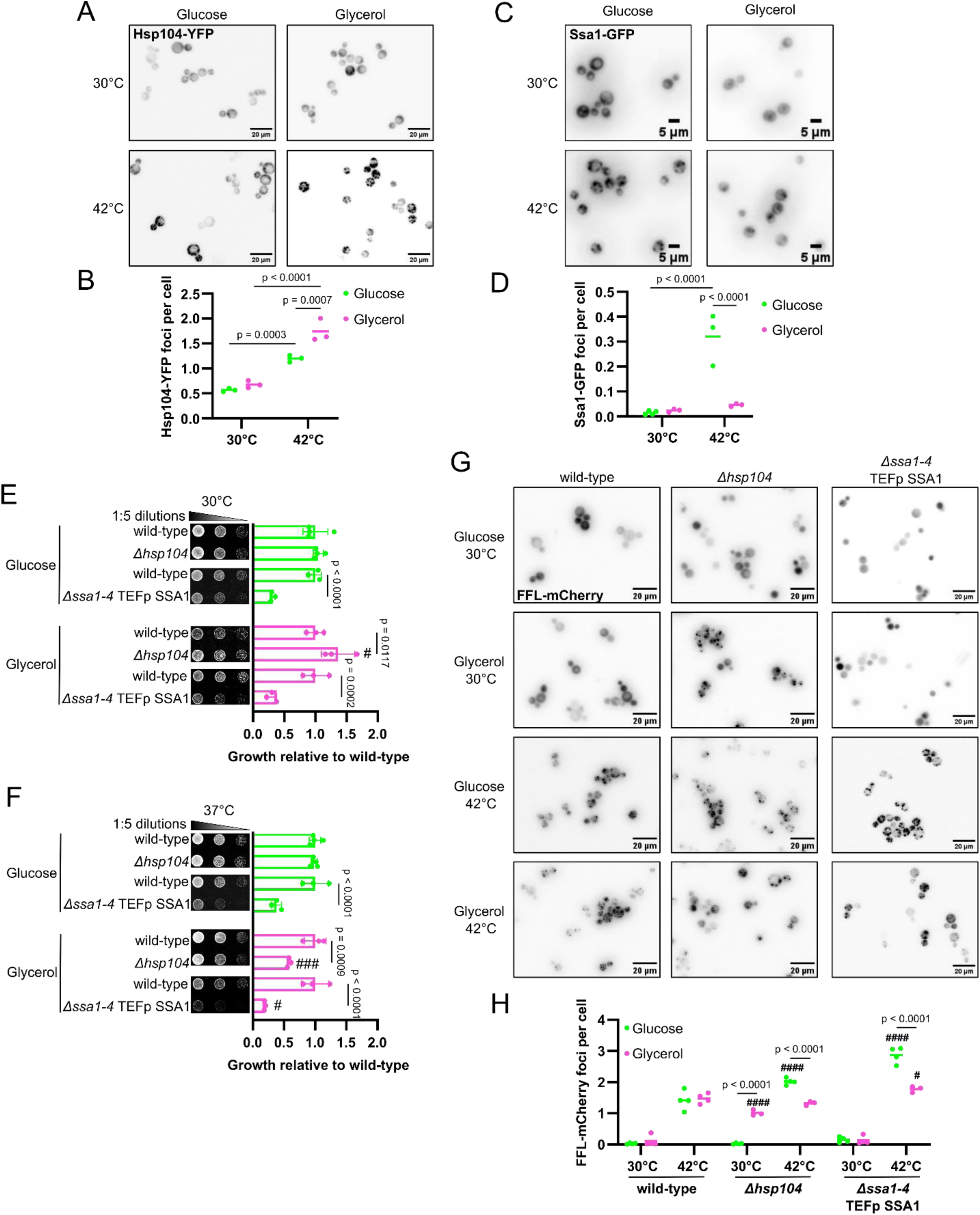
Reprogramming of Hsp104 and Hsp70 in respiring cells. A. Fluorescence microscopy and B. quantification of the number of Hsp104-YFP foci per cell in cells grown in media supplemented with either glucose or glycerol at 30°C or heat shocked at 42°C for 1 hour. C. Fluorescence microscopy and D. quantification of the number of Ssa1-GFP foci per cell in cells grown in media supplemented with either glucose or glycerol at 30°C or heat shocked at 42°C for 1 hour. Growth assays of wild-type, *Δhsp104*, and *Δssa1-4* TEFp SSA1 cells grown on plates supplemented with either glucose or glycerol at E. 30°C or F. 37°C. The mean growth (+/- SD) relative wild-type cells are represented graphically for at least three independent experiments. # symbols indicate significant differences compared to the same sample metabolizing glucose at the same temperature (*# p<0.05, ## p<0.01, ### p<0.001, #### p<0.0001*). G. Fluorescence microscopy and H. quantification of the number of FFL-mCherry foci per cell in wild-type, *Δhsp104*, and *Δssa1-4* TEFp SSA1 cells grown in media supplemented with either glucose or glycerol at 30°C or heat shocked at 42°C for 1 hour. The number of FFL-mCherry foci per cell is represented graphically for at least three independent experiments. # symbols indicate significant differences compared to wild-type cells metabolizing glucose at the same temperature (*# p<0.05, ## p<0.01, ### p<0.001, #### p<0.0001*).

We next explored whether Hsp104 and Hsp70 regulate the viability of respiring cells. To this end, we performed growth assays using cells deleted for HSP104 (*Δhsp104*) or expressing reduced cytosolic Hsp70 levels (*Δssa1-4* TEFp SSA1) [15,16]. We find that *Δhsp104* cells grow at a similar rate to wild-type cells, while *Δssa1-4* TEFp SSA1 cells exhibit a growth defect when metabolizing glucose at 30°C (Fig. 5E). Interestingly, *Δhsp104* cells show improved growth, while *Δssa1-4* TEFp SSA1 cells show no difference in growth behavior when metabolizing glycerol at 30°C (Fig. 5E). Growth at 37°C does not change the growth of *Δhsp104* or *Δssa1-4* TEFp SSA1 cells metabolizing glucose (Fig. 5F). However, both *Δhsp104* and *Δssa1-4* TEFp SSA1 cells are sensitive to elevated temperature when metabolizing glycerol (Fig. 5F).

We then turned our attention to whether Hsp104 and Hsp70 regulate proteostasis of respiring cells. Accordingly, we monitored aggregation of the proteostasis reporter FFL-mCherry in *Δhsp104* and *Δssa1-4* TEFp SSA1 cells. FFL-mCherry is soluble in wild-type, *Δhsp104* and *Δssa1-4* TEFp SSA1 cells grown in media containing glucose at 30°C (Fig. 5G-H). Metabolism of glycerol at 30°C only induces an increase in FFL-mCherry aggregation in *Δhsp104* cells, but not wild-type or *Δssa1-4* TEFp SSA1 cells (Fig. 5G-H). A 1-hour heat shock of wild-type, *Δhsp104* and *Δssa1-4* TEFp SSA1 cells metabolizing glucose increases FFL-mCherry aggregation when compared to cells metabolizing glucose at 30°C (Fig. 5G-H). Notably, the number of FFL-mCherry aggregates following a heat shock is increased in *Δhsp104* and *Δssa1-4* TEFp SSA1 cells metabolizing glucose when compared to wild-type cells (Fig. 5G-H). Interestingly, a heat shock induces fewer FFL-mCherry aggregates in *Δhsp104* and *Δssa1-4* TEFp SSA1 cells metabolizing glycerol when compared to cells metabolizing glucose (Fig. 5G-H).

In sum, the expression and functions of Hsp104 and Hsp70 in response to stress are modified in respiring cells. Of particular note, Hsp104 is overemployed to inclusions in stressed respiring cells, while the ability of Ssa1 to localize to inclusions is abolished in respiring cells. In addition, we find that both Hsp104 and Hsp70 are more critical for viability and proteostasis in respiring cells than in glycolytic cells.

## DISCUSSION

When protein folding is challenged, cells rapidly mount transcriptional responses that promote the expression of molecular chaperones and protein remodeling complexes to restore proteostasis [1–4]. The heat shock response is specifically employed in response to nuclear and cytosolic protein misfolding and is regulated by activation of Hsf1 [1,7–10,24]. While the role of the heat shock response in tolerance of protein misfolding and aggregation is well understood in glycolytic cells, it is currently unknown whether the heat shock response functions identically in respiring cells, such as neurons. Here, we manipulate the metabolism of yeast by changing the carbon source available in the media to determine how proteostasis is mediated by the heat shock response in respiring cells. Although this model allows for simple manipulation of energy metabolism, it also promotes changes in growth rate. Thus, it is possible, though unlikely in the time scales that we utilize, that slower rates of cell division in respiring cells contribute to impaired proteostasis we observe [77].

We validate that changing the carbon source available in the media from glucose to glycerol induces transcriptional changes that downregulate genes involved in glycolytic metabolism and upregulates genes involved in oxidative phosphorylation, indicating a shift in metabolic program. We find that, although both glycolytic and respiring cells upregulated transcripts associated with the heat shock response when stressed, respiring cells were deficient in upregulation of these transcripts. The impaired heat shock response in respiring cells is also associated with an inability to trigger the condensation cascade required for Hsf1 activity. A such, we note reduced gene expression and protein steady-state levels of the Hsf1 regulated molecular chaperones Hsp104, Hsp70 and Hsp26 in respiring cells. Notably, we identified an increase in transcript levels of genes associated with ribosome biogenesis in respiring cells when stressed, which corresponds with a lack of protein biosynthesis depression associated with heat shock [66]. Interestingly, the maintained protein biosynthesis in respiring cells corresponds to a lack of oRPs that typically initiate Hsf1 activation [67,69,70]. Thus, we speculate that carbon source regulates the translational response to heat shock upstream of Hsf1 activation to dampen the heat shock response.

In this study, we explore how the impaired heat shock response influences protein misfolding and aggregation in respiring cells. Corresponding to changes in molecular chaperone expression, we find changes in spatial protein quality control in respiring cells. Specifically, respiring cells are less efficient at dispersing aggregates of metastable proteins. These findings parallel our previous findings that metabolism modifies aggregation of neurodegenerative disease-associated proteins TDP-43 and FUS [52]. Triaging of misfolded proteins into protein quality control compartments facilitates their sequestration and degradation, plausibly modifying their toxicity [73,78–80]. While we focus on characterizing the heat shock response and changes in molecular chaperone function in respiring cells, our work highlights an additional need for characterizing differences in the formation and function of spatial protein quality control compartments in respiring cells.

Hsf1 is often described as a pro-survival transcription factor due to its association with tolerance of stress [20–22,81]. Specifically, loss of Hsf1 activity is associated with proteostasis decline in ageing cells [14,82–86], which can be overcome by artificial induction of the heat shock response [23]. However, overactivation of the heat shock response can be detrimental [87], illustrating the importance of stringent regulation of Hsf1. Interestingly, we demonstrate that impaired activity of Hsf1 is adaptive in respiring cells, as an overactive Hsf1 variant is detrimental to the growth of respiring cells while an inhibited Hsf1 variant is well tolerated in respiring cells. Additionally, although impaired Hsf1 function unsurprisingly impairs proteostasis in glycolytic cells, we demonstrate that impaired Hsf1 function instead reduces aggregation of metastable proteins in respiring cells. Because inhibition of Hsf1 is crucial to the viability of respiring cells, we speculate that instead of regulating proteostasis by increasing the pool of molecular chaperones available for protein folding, respiring cells regulate proteostasis by modifying the activities of pre-existing molecular chaperones.

Clearance of protein aggregates is regulated by the protein remodeling factor Hsp104 and the molecular chaperone Hsp70 [12,16,74–76]. As we identified that respiring cells are inefficient at clearing protein aggregates following stress, we hypothesized that the functions of Hsp104 and Hsp70, in addition to their expression, are dysregulated in respiring cells. In accordance with their functions in protein aggregate clearance, both Hsp104 and Hsp70 are localized to cytosolic inclusions in stressed glycolytic cells. However, we demonstrate partitioning of their functions in respiring cells. Hsp104 is overemployed into cytosolic inclusions in response to stress, while Hsp70 remains diffusely localized. We speculate that respiring cells employ the more abundant Hsp70 to mitigate the misfolding of soluble proteins. Thus, only Hsp104 remains to mitigate protein aggregation, preventing efficient aggregate clearance. Accordingly, loss of function of Hsp104 induces protein aggregation in respiring cells in the absence of stress. Furthermore, loss of function of either Hsp104 or Hsp70 is only detrimental to stressed cells growing at elevated temperatures, demonstrating a specific demand for these molecular chaperones in respiring cells responding to preteotostatic stress. Taken together, our findings demonstrate a genetic mechanism by which respiring cells reprogram molecular chaperones to perform distinct functions to respond to stress-induced proteostatic burden.

## CONCLUSION

Energy demanding cells, such as neurons, employ oxidative phosphorylation to generate ATP efficiently at the expense of redox homeostasis. Yet, whether oxidative phosphorylation challenges the maintenance of proteostasis independent of oxidative stress was hitherto underexplored. Here, we demonstrate that altering the metabolic program from glycolysis to oxidative phosphorylation impairs Hsf1 activation, the heat shock response, and in turn, the maintenance of cytosolic proteostasis. Remarkably, artificially increasing the heat shock response in respiring cells is detrimental and does not improve proteostasis. Instead, our data indicate that respiring cells redirect the existing pool of molecular chaperones towards distinct functions to mitigate protein misfolding in response to stress. Our findings provide a compelling genetic model whereby Hsp70 is redirected away from protein aggregates toward refolding soluble misfolded proteins, leaving Hsp104 as an independent regulator of protein disaggregation in respiring cells. Taken together, our findings suggest that metabolic program compounds the existing nuance in proteostasis regulation. In addition, we provide a plausible mechanism by which respiring metazoan cells, including neurons, may be specifically susceptible to protein misfolding associated with ageing and neurodegenerative diseases. Finally, we demonstrate a need to consider the impact of distinct metabolic activities when exploring protein misfolding diseases, including neurodegenerative diseases.

## MATERIALS & METHODS

### Yeast strains, plasmids and growth conditions

All yeast strains used in this study are derivatives of either W303 or BY4742 and are described in Table S1. The three HSF1 mutant strains (HSF1 wt, *hsf1ΔN*, and *hsf1ΔC*) were described previously [8]. The *Δhsp104* strain was described previously [16]. The strain with reduced expression of cytosolic Hsp70 (*Δssa1-4* TEFp SSA1) was kindly gifted to us by Dr. Elizabeth Craig [15]. The SSA1-GFP strain was acquired from the yeast GFP clone collection [88].

For ectopic protein expression, plasmids were transformed into yeast cells using the standard LiAc/PEG protocol [89]. The proteostasis reporter plasmid, p426MET17 FFL_D50N,G119N_-GFP was a kind gift from Dr. John Glover [53]. A low copy number p416GPD FFL_D50N,G119N_-mCherry was generated by cut paste cloning. The p415GPD HSP104-YFP plasmid used to monitor Hsp104 inclusion formation was described previously.

To induce metabolism by glycolysis, cells were grown in liquid or plated on agar plates containing synthetic defined (SD) media supplemented with 2% glucose, 6.7 g/mL yeast nitrogenous bases with ammonium sulfate and amino acids (40 mg/mL L-lysine, 20 mg/mL L-arginine, 10 mg/mL L-threonine, 60 mg/mL L-phenylalanine, 20 mg/mL L-isoleucine, 10 mg/mL L-methionine, 20 mg/mL adenine hemisulfate). For selection, the media was supplemented with additional amino acids (20 mg/mL L-histidine, 60 mg/mL L-leucine, 20 mg/mL uracil, 80 mg/mL L-tryptophan). To induce metabolism by oxidative phosphorylation, 2% glucose was replaced with 2% glycerol or 2% potassium acetate in the media. To induce metabolism by β-oxidation, 2% glucose was replaced with 0.1% myristic acid or 0.1% oleic acid, both of which were first solubilized in Tween40 (final concentration of 0.05%).

For HaloTag imaging, cells were grown to mid-log phase at 30°C in YPD (1% yeast extract, 2% peptone, 2% glucose) or YP-glycerol (1% yeast extract, 2% peptone, 2% glycerol).

### RNA extraction and sequencing

RNA was extracted from three independent biological replicates using TRIzol extraction. Briefly, five OD600 units of mid-log phase cells were pelleted and resuspended in 100 µL TRIzol and transferred to an eppendorf tube containing an equal volume of acid-washed glass beads. The cells were then disrupted using the Analog Disruptor Genie (Scientific Industries, SI-D238) for three 60-second intervals interspersed by 60-second incubations on ice. The lysate was then transferred to a new tube, diluted with 1 mL of TRIzol and treated with 200 µL chloroform for 5 minutes at room temperature. The lysate was then centrifuged at 12k xg for 15 minutes at 4°C, and the aqueous phase was transferred to a new eppendorf tube. The RNA was then purified using the Purelink RNA Mini Kit (ThermoFisher, 12183018A) as per manufacturer instructions.

Total RNA was submitted to GenomeQuébec for sequencing, where the samples were depleted of rRNA, and cDNA libraries were prepared and sequenced using Illumina NovaSeq X. 26-46 million raw reads were received for each sample. An alignment index was prepared for R64 strain S288C (GenBank ID: GCF_000146045.2) using STARv2.7.11b through the RSEMv1.3.3 tool [90,91]. STAR’s splice junction database overhang parameter was set to 99 nucleotides. The raw FASTQ reads were aligned to the R64 genome with STAR through the RSEM tool. Reverse strandedness was specified. Cohort-wide count tables were created by combining each sample’s independent expression files. Additional quality control was performed on the raw counts to identify potential outliers and sample biases. Differentially expressed genes (DEG) were assessed in R using DESeq2 [92]. These data are available upon request.

### Transcriptome analysis

Principal component analysis was performed on the cohort-wide normalized count tables using R. Volcano plots were generated using differential expression analysis (DEA) tables for each comparison using R. Enrichment analysis of gene ontology (GO) terms and transcription factors (TFs) in sets of upregulated and downregulated genes was assessed using g:Profiler [93]. The full list of enriched GO terms and TFs for each comparison is available in Table S2-7). Dotplots highlighting enriched GO terms and TFs were generated using R. All R scripts used for transcriptome analysis in this study are available at https://doi.org/10.6084/m9.figshare.32207130.

### Pulse-labelling of nascent peptides

Nascent polypeptides were labelled using click chemistry as described previously [64,65]. To label nascent polypeptides, liquid cell cultures were resuspended in media lacking methionine and containing 50 µM L-Azidohomoalanine (AHA) for five minutes prior to or following a 1-hour heat shock at 42°C. AHA labelled peptides were then labelled with 10 µM sDIBO alkyne AlexaFluor488 overnight at room temperature. Labelling was repeated using samples that were not treated with AHA as a control. Lysates were then resolved using a stain-free SDS-PAGE (Bio-Rad, 4568023). Labelled proteins were imaged using ChemiDoc MP (Bio-Rad) and densitometry of was measured using Fiji [94].

### HaloTag pulse labelling and multi-channel fluorescence microscopy

To assess orphan ribosome proteins, Rpl26a-HaloTag was pulse-labeled with JF-646 HaloTag ligand in cells heat shocked at 39°C for 15 minutes prior to fixation as described [67]. Cells were fixed in 4% paraformaldehyde and imaged as described [68,95]. Multi-channel fluorescence microscopy images were acquired on a Nikon SoRa spinning disk confocal microscope (63× objective) at the University of Chicago Integrated Light Microscopy Core (RRID: SCR_019197). Images were collected as z-stacks and converted to maximum intensity projections prior to analysis. Image analysis was performed using cellquant (version 1.0) [96]. Cell segmentation used the Cellpose cpsam model with composite-channel segmentation input and cell area filtering (200–5,000 pixels). Colocalization was computed as Pearson’s r with Costes automatic thresholding. The condensate index was defined as the ratio of the 95th percentile to the mean pixel intensity within each segmented cell, providing a measure of signal concentration into [96]. Per-cell measurements were summarized at the replicate level using medians. T-tests were performed on replicate-level medians. n ≥ 3 biological replicates per condition, with >200 cells per replicate. Only comparisons reaching p < 0.05 are annotated. Data are presented as superplots in which individual cells are shown as small dots and replicate medians as large diamonds.

### Growth assay

Growth assays to determine growth defects were performed as described in Petropavlovskiy et al. [97]. Briefly, cells were diluted to OD600=1 in a 96-well and serial 5-fold dilutions were performed. Cells were then plated onto agar plates using a 48-pronged frogger and incubated at either 30°C or 37°C for 24-72 hours. Plates were then imaged and the densitometry of the spots was analyzed using Fiji [94]. Each experiment was performed at least three times using independent replicates.

### Single channel fluorescence microscopy

To measure aggregation or inclusion formation, we performed fluorescence microscopy. Cells were grown to mid-log phase prior to microscopy. For assessment of proteostasis with the FFL-GFP reporter, FFL-GFP was induced for 1 hour by resuspending cells in media lacking methionine. Where indicated, cells were subjected to a 1-hour heat shock at 42°C. Cells were then imaged with the BioTek Cytation 5 Cell Imaging Multi-mode Reader at 20x magnification using the GFP and Texas-Red filters where appropriate. For cells grown in myristic acid or oleic acid media, cells were plated onto selective agar plates supplemented with myristic acid and oleic acid and lacking methionine to induce the expression of FFL-GFP. After two days of growth, cells were scraped off the plate and immediately imaged. Microscopy was repeated with at least three independent replicates and at least 500 cells were imaged for each condition. Images were analyzed for foci formation using the EBImage library in R [98]. The R script for quantification of micrographs used in this study is available at https://doi.org/10.6084/m9.figshare.28688966.

### Protein extraction and western blot

Cells were grown to mid-log phase and lysed following a standard alkaline lysis protocol [99]. Briefly, one OD600 unit of mid-log cells were pelleted and resuspended in 100 mM NaOH, 2% SDS, 50 mM EDTA and 2% 2-mercaptoethanol, boiled for five minutes, diluted in an equal volume of 1% SDS, 8 M Urea, 10 mM MOPS pH 6.8, 10 mM EDTA and 0.01% bromophenol blue and boiled again for five minutes. Proteins were then resolved via SDS-PAGE and transferred to a nitrocellulose membrane. The membranes were then blocked with 5% skim milk powder in TBST before incubation with primary antibody diluted in 1% BSA in TBST overnight at 4C. The primary antibodies used in this study are: FFL (Rb anti-FFL, 1:1000, abcam ab21176), Hsp104 (Rb anti-Hsp104, 1:5000, gift from Dr. Johannes Buchner), Hsp70 (Ms anti-Hsp70 1:1000, Thermofisher MA3-006) and the loading control Pgk1 (Rb anti-Pgk1 1:5000, Origene AP21371AF-N). Subsequently, membranes were washed with TBST before incubation with secondary antibody diluted 1:5000 in 1% BSA in TBST for one hour at room temperature. The secondary antibodies used in this study are: Gt anti Rb IgG AlexaFluor680 (Thermofisher, A32734) and Dk anti Ms IgG AlexaFluor680 (Thermofisher, A10038). Membranes were then imaged using the ChemiDoc MP (Bio-Rad) and densitometry of was measured using Fiji [94].

## ACKNOWLEDGEMENTS

The authors would like to thank Drs. Elizabeth Craig (Emeritus, University of Wisconsin-Madison) and John Glover (University of Toronto) for graciously providing strains and plasmids used in this study. The authors acknowledge the University of Chicago Integrated Light Microscopy Core (RRID: SCR_019197) for providing equipment and services used in this study. We would like to thank Drs. Patrick Lajoie (University of Western Ontario), Paul Walton (Emeritus, University of Western Ontario), Jim Karagiannis (University of Western Ontario), and Sue-Ann Mok (University of Alberta) for their critical reading of this manuscript. This research was funded through a NSERC Discovery grant (RGPIN-2024-05867) and a McGill-Western Initiative for Translational Neuroscience (ITN) grant to M.L.D., an ALS Canada Trainee Fellowship to D.W.M, National Institutes of Health grants R01 GM138689 and RM1 GM153533 and National Science Foundation QLCI QuBBE grant OMA-2121044 to D.P. This research was also supported by CIHR Skin Research Training Centre (201903).

## REFERENCES

1. Morimoto RI, Santoro MG. Stress–inducible responses and heat shock proteins: New pharmacologic targets for cytoprotection. Nat Biotechnol. 1998;16: 833–838. doi:10.1038/nbt0998-833

2. Hipp MS, Kasturi P, Hartl FU. The proteostasis network and its decline in ageing. Nat Rev Mol Cell Biol. 2019;20: 421–435. doi:10.1038/s41580-019-0101-y

3. Hetz C, Zhang K, Kaufman RJ. Mechanisms, regulation and functions of the unfolded protein response. Nat Rev Mol Cell Biol. 2020;21: 421–438. doi:10.1038/s41580-020-0250-z

4. Hartl FU. Molecular chaperones in cellular protein folding. Nature. 1996;381: 571–9. doi:10.1038/381571a0

5. Sorger PK, Pelham HR. Purification and characterization of a heat-shock element binding protein from yeast. EMBO J. 1987;6: 3035–3041. doi:10.1002/j.1460-2075.1987.tb02609.x

6. Jakobsen BK, Pelham HRB. Constitutive Binding of Yeast Heat Shock Factor to DNA In Vivo. Mol Cell Biol. 1988;8: 5040–5042. doi:10.1128/mcb.8.11.5040-5042.1988

7. Zheng X, Krakowiak J, Patel N, Beyzavi A, Ezike J, Khalil AS, et al. Dynamic control of Hsf1 during heat shock by a chaperone switch and phosphorylation. Hunter T, editor. Elife. 2016;5: e18638. doi:10.7554/eLife.18638

8. Krakowiak J, Zheng X, Patel N, Feder ZA, Anandhakumar J, Valerius K, et al. Hsf1 and Hsp70 constitute a two-component feedback loop that regulates the yeast heat shock response. Hunter T, editor. Elife. 2018;7: e31668. doi:10.7554/eLife.31668

9. Sorger PK, Pelham HRB. Yeast heat shock factor is an essential DNA-binding protein that exhibits temperature-dependent phosphorylation. Cell. 1988;54: 855–864. doi:10.1016/S0092-8674(88)91219-6

10. Trotter EW, Kao CM-F, Berenfeld L, Botstein D, Petsko GA, Gray J V. Misfolded Proteins Are Competent to Mediate a Subset of the Responses to Heat Shock in Saccharomyces cerevisiae *. Journal of Biological Chemistry. 2002;277: 44817–44825. 10.1074/jbc.M204686200

11. Zhang H, Amick J, Chakravarti R, Santarriaga S, Schlanger S, McGlone C, et al. A Bipartite Interaction between Hsp70 and CHIP Regulates Ubiquitination of Chaperoned Client Proteins. Structure. 2015;23: 472–482. doi:10.1016/j.str.2015.01.003

12. Yoo H, Bard JAM, Pilipenko E V, Drummond DA. Chaperones directly and efficiently disperse stress-triggered biomolecular condensates. Mol Cell. 2022;82: 741–755.e11. doi:10.1016/j.molcel.2022.01.005

13. Hanzén S, Vielfort K, Yang J, Roger F, Andersson V, Zamarbide-Forés S, et al. Lifespan Control by Redox-Dependent Recruitment of Chaperones to Misfolded Proteins. Cell. 2016;166: 140–151. doi:10.1016/j.cell.2016.05.006

14. Morley JF, Morimoto RI. Regulation of Longevity in *Caenorhabditis elegans* by Heat Shock Factor and Molecular Chaperones. Mol Biol Cell. 2004;15: 657–664. doi:10.1091/mbc.e03-07-0532

15. Schilke BA, Ciesielski SJ, Ziegelhoffer T, Kamiya E, Tonelli M, Lee W, et al. Broadening the functionality of a J-protein/Hsp70 molecular chaperone system. PLoS Genet. 2017;13: e1007084. doi:10.1371/journal.pgen.1007084

16. Cashikar AG, Duennwald M, Lindquist SL. A Chaperone Pathway in Protein Disaggregation: HSP26 ALTERS THE NATURE OF PROTEIN AGGREGATES TO FACILITATE REACTIVATION BY HSP104*. Journal of Biological Chemistry. 2005;280: 23869–23875. 10.1074/jbc.M502854200

17. Freeman BC, Morimoto RI. The human cytosolic molecular chaperones hsp90, hsp70 (hsc70) and hdj-1 have distinct roles in recognition of a non-native protein and protein refolding. EMBO J. 1996;15: 2969–79.

18. Quintana-Gallardo L, Martín-Benito J, Marcilla M, Espadas G, Sabidó E, Valpuesta JM. The cochaperone CHIP marks Hsp70- and Hsp90-bound substrates for degradation through a very flexible mechanism. Sci Rep. 2019;9: 5102. doi:10.1038/s41598-019-41060-0

19. Xiao X. HSF1 is required for extra-embryonic development, postnatal growth and protection during inflammatory responses in mice. EMBO J. 1999;18: 5943–5952. doi:10.1093/emboj/18.21.5943

20. Dai C, Whitesell L, Rogers AB, Lindquist S. Heat Shock Factor 1 Is a Powerful Multifaceted Modifier of Carcinogenesis. Cell. 2007;130: 1005–1018. doi:10.1016/j.cell.2007.07.020

21. Jolly C. Role of the Heat Shock Response and Molecular Chaperones in Oncogenesis and Cell Death. J Natl Cancer Inst. 2000;92: 1564–1572. doi:10.1093/jnci/92.19.1564

22. Zhou Z, Li Y, Jia Q, Wang Z, Wang X, Hu J, et al. Heat shock transcription factor 1 promotes the proliferation, migration and invasion of osteosarcoma cells. Cell Prolif. 2017;50. doi:10.1111/cpr.12346

23. Harris N, MacLean M, Hatzianthis K, Panaretou B, Piper PW. Increasing Saccharomyces cerevisiae stress resistance, through the overactivation of the heat shock response resulting from defects in the Hsp90 chaperone, does not extend replicative life span but can be associated with slower chronological ageing of nondividing cells. Molecular Genetics and Genomics. 2001;265: 258–263. doi:10.1007/s004380000409

24. Solís EJ, Pandey JP, Zheng X, Jin DX, Gupta PB, Airoldi EM, et al. Defining the Essential Function of Yeast Hsf1 Reveals a Compact Transcriptional Program for Maintaining Eukaryotic Proteostasis. Mol Cell. 2016;63: 60–71. 10.1016/j.molcel.2016.05.014

25. Klaips CL, Gropp MHM, Hipp MS, Hartl FU. Sis1 potentiates the stress response to protein aggregation and elevated temperature. Nat Commun. 2020;11: 6271. doi:10.1038/s41467-020-20000-x

26. Park S-K, Hong JY, Arslan F, Kanneganti V, Patel B, Tietsort A, et al. Overexpression of the essential Sis1 chaperone reduces TDP-43 effects on toxicity and proteolysis. PLoS Genet. 2017;13: e1006805. doi:10.1371/journal.pgen.1006805

27. Park S-K, Arslan F, Kanneganti V, Barmada SJ, Purushothaman P, Verma SC, et al. Overexpression of a conserved HSP40 chaperone reduces toxicity of several neurodegenerative disease proteins. Prion. 2018;12: 16–22. doi:10.1080/19336896.2017.1423185

28. Lin LTW, Razzaq A, Di Gregorio SE, Hong S, Charles B, Lopes MH, et al. Hsp90 and its co-chaperone Sti1 control TDP-43 misfolding and toxicity. The FASEB Journal. 2021;35: e21594. doi:10.1096/FJ.202002645R

29. Warburg O. □ber den Stoffwechsel der Carcinomzelle. Naturwissenschaften. 1924;12: 1131–1137. doi:10.1007/BF01504608

30. Warburg O. On the Origin of Cancer Cells. Science (1979). 1956;123: 309–314. doi:10.1126/science.123.3191.309

31. Mitchell P. Coupling of Phosphorylation to Electron and Hydrogen Transfer by a Chemi-Osmotic type of Mechanism. Nature. 1961;191: 144–148. doi:10.1038/191144a0

32. Guppy M, Greiner E, Brand K. The role of the Crabtree effect and an endogenous fuel in the energy metabolism of resting and proliferating thymocytes. Eur J Biochem. 1993;212: 95–99. doi:10.1111/j.1432-1033.1993.tb17637.x

33. Pfeiffer T, Schuster S, Bonhoeffer S. Cooperation and Competition in the Evolution of ATP-Producing Pathways. Science (1979). 2001;292: 504–507. doi:10.1126/science.1058079

34. Agathocleous M, Love NK, Randlett O, Harris JJ, Liu J, Murray AJ, et al. Metabolic differentiation in the embryonic retina. Nat Cell Biol. 2012;14: 859–864. doi:10.1038/ncb2531

35. Zheng X, Boyer L, Jin M, Mertens J, Kim Y, Ma L, et al. Metabolic reprogramming during neuronal differentiation from aerobic glycolysis to neuronal oxidative phosphorylation. Elife. 2016;5. doi:10.7554/eLife.13374

36. Schurr A, Payne RS. Lactate, not pyruvate, is neuronal aerobic glycolysis end product: An in vitro electrophysiological study. Neuroscience. 2007;147: 613–619. doi:10.1016/j.neuroscience.2007.05.002

37. Ross JM, Öberg J, Brené S, Coppotelli G, Terzioglu M, Pernold K, et al. High brain lactate is a hallmark of aging and caused by a shift in the lactate dehydrogenase A/B ratio. Proceedings of the National Academy of Sciences. 2010;107: 20087–20092. doi:10.1073/pnas.1008189107

38. Poderoso JJ, Carreras MC, Lisdero C, Riobó N, Schöpfer F, Boveris A. Nitric Oxide Inhibits Electron Transfer and Increases Superoxide Radical Production in Rat Heart Mitochondria and Submitochondrial Particles. Arch Biochem Biophys. 1996;328: 85–92. doi:10.1006/abbi.1996.0146

39. Loschen G, Flohé L. Respiratory chain linked H _2_ O _2_ production in pigeon heart mitochondria. FEBS Lett. 1971;18: 261–264. doi:10.1016/0014-5793(71)80459-3

40. Sridharan S, Kurzawa N, Werner T, Günthner I, Helm D, Huber W, et al. Proteome-wide solubility and thermal stability profiling reveals distinct regulatory roles for ATP. Nat Commun. 2019;10: 1155. doi:10.1038/s41467-019-09107-y

41. Patel A, Malinovska L, Saha S, Wang J, Alberti S, Krishnan Y, et al. ATP as a biological hydrotrope. Science (1979). 2017;356: 753–756. doi:10.1126/science.aaf6846

42. Hayes MH, Peuchen EH, Dovichi NJ, Weeks DL. Dual roles for ATP in the regulation of phase separated protein aggregates in Xenopus oocyte nucleoli. Elife. 2018;7. doi:10.7554/eLife.35224

43. Takaine M, Imamura H, Yoshida S. High and stable ATP levels prevent aberrant intracellular protein aggregation in yeast. Elife. 2022;11. doi:10.7554/eLife.67659

44. Carlson M. Glucose repression in yeast. Curr Opin Microbiol. 1999;2: 202–207. doi:10.1016/S1369-5274(99)80035-6

45. Herrero P, Fern□ndez R, Moreno F. Differential sensitivities to glucose and galactose repression of gluconeogenic and respiratory enzymes from Saccharomyces cerevisiae. Arch Microbiol. 1985;143: 216–219. doi:10.1007/BF00411238

46. Schüller H-J. Transcriptional control of nonfermentative metabolism in the yeast Saccharomyces cerevisiae. Curr Genet. 2003;43: 139–160. doi:10.1007/s00294-003-0381-8

47. Burtner CR, Murakami CJ, Kennedy BK, Kaeberlein M. A molecular mechanism of chronological aging in yeast. Cell Cycle. 2009;8: 1256–70. doi:10.4161/cc.8.8.8287

48. Sokolov S, Pozniakovsky A, Bocharova N, Knorre D, Severin F. Expression of an expanded polyglutamine domain in yeast causes death with apoptotic markers. Biochimica et Biophysica Acta (BBA) - Bioenergetics. 2006;1757: 660–666. doi:10.1016/j.bbabio.2006.05.004

49. Braun RJ, Sommer C, Carmona-Gutierrez D, Khoury CM, Ring J, Büttner S, et al. Neurotoxic 43-kDa TAR DNA-binding Protein (TDP-43) Triggers Mitochondrion-dependent Programmed Cell Death in Yeast. Journal of Biological Chemistry. 2011;286: 19958–19972. doi:10.1074/jbc.M110.194852

50. Ocampo A, Zambrano A, Barrientos A. Suppression of polyglutamine-induced cytotoxicity in *Saccharomyces cerevisiae* by enhancement of mitochondrial biogenesis. The FASEB Journal. 2010;24: 1431–1441. doi:10.1096/fj.09-148601

51. Akintade DD, Chaudhuri B. Apoptosis, Induced by Human α-Synuclein in Yeast, Can Occur Independent of Functional Mitochondria. Cells. 2020;9. doi:10.3390/cells9102203

52. McDonald DW, Chugh N, Sava R, Duennwald ML. FUS and TDP-43 aggregation are uncoupled from toxicity in ageing yeast models. BMC Biol. 2026. doi:10.1186/s12915-026-02537-3

53. Tkach JM, Glover JR. Nucleocytoplasmic Trafficking of the Molecular Chaperone Hsp104 in Unstressed and Heat-Shocked Cells. Traffic. 2008;9: 39–56. doi:10.1111/j.1600-0854.2007.00666.x

54. Strich R, Surosky RT, Steber C, Dubois E, Messenguy F, Esposito RE. UME6 is a key regulator of nitrogen repression and meiotic development. Genes Dev. 1994;8: 796–810. doi:10.1101/gad.8.7.796

55. Park H-D, Luche RM, Cooper TG. The yeast *UME6* gene product is required for transcriptional repression mediated by the *CAR1 URS1* repressor binding site. Nucleic Acids Res. 1992;20: 1909–1915. doi:10.1093/nar/20.8.1909

56. Bojunga N, Entian KD. Cat8p, the activator of gluconeogenic genes in Saccharomyces cerevisiae, regulates carbon source-dependent expression of NADP-dependent cytosolic isocitrate dehydrogenase (Idp2p) and lactate permease (Jen1p). Mol Gen Genet. 1999;262: 869–75. doi:10.1007/s004380051152

57. Hedges D, Proft M, Entian K-D. *CAT8*, a New Zinc Cluster-Encoding Gene Necessary for Derepression of Gluconeogenic Enzymes in the Yeast *Saccharomyces cerevisiae*. Mol Cell Biol. 1995;15: 1915–1922. doi:10.1128/MCB.15.4.1915

58. Mai B, Breeden L. Xbp1, a stress-induced transcriptional repressor of the Saccharomyces cerevisiae Swi4/Mbp1 family. Mol Cell Biol. 1997;17: 6491–501. doi:10.1128/MCB.17.11.6491

59. Miles S, Li L, Davison J, Breeden LL. Xbp1 directs global repression of budding yeast transcription during the transition to quiescence and is important for the longevity and reversibility of the quiescent state. PLoS Genet. 2013;9: e1003854. doi:10.1371/journal.pgen.1003854

60. Ozcan S, Leong T, Johnston M. Rgt1p of Saccharomyces cerevisiae, a key regulator of glucose-induced genes, is both an activator and a repressor of transcription. Mol Cell Biol. 1996;16: 6419–26. doi:10.1128/MCB.16.11.6419

61. Marshall-Carlson L, Neigeborn L, Coons D, Bisson L, Carlson M. Dominant and recessive suppressors that restore glucose transport in a yeast snf3 mutant. Genetics. 1991;128: 505–12. doi:10.1093/genetics/128.3.505

62. Cairns BR, Lorch Y, Li Y, Zhang M, Lacomis L, Erdjument-Bromage H, et al. RSC, an essential, abundant chromatin-remodeling complex. Cell. 1996;87: 1249–60. doi:10.1016/s0092-8674(00)81820-6

63. Angus-Hill ML, Schlichter A, Roberts D, Erdjument-Bromage H, Tempst P, Cairns BR. A Rsc3/Rsc30 zinc cluster dimer reveals novel roles for the chromatin remodeler RSC in gene expression and cell cycle control. Mol Cell. 2001;7: 741–51. doi:10.1016/s1097-2765(01)00219-2

64. Morey TM, Esmaeili MA, Duennwald ML, Rylett RJ. SPAAC Pulse-Chase: A Novel Click Chemistry-Based Method to Determine the Half-Life of Cellular Proteins. Front Cell Dev Biol. 2021;9. doi:10.3389/FCELL.2021.722560

65. McDonald DW, Dib RN, De Luca C, Shah A, Duennwald ML. Specific branches of the proteostasis network regulate the toxicity associated with mistranslation. Nucleic Acids Res. 2025;53. doi:10.1093/nar/gkaf428

66. Mühlhofer M, Berchtold E, Stratil CG, Csaba G, Kunold E, Bach NC, et al. The Heat Shock Response in Yeast Maintains Protein Homeostasis by Chaperoning and Replenishing Proteins. Cell Rep. 2019;29: 4593–4607.e8. doi:10.1016/j.celrep.2019.11.109

67. Ali A, Garde R, Schaffer OC, Bard JAM, Husain K, Kik SK, et al. Adaptive preservation of orphan ribosomal proteins in chaperone-dispersed condensates. Nat Cell Biol. 2023;25: 1691–1703. doi:10.1038/s41556-023-01253-2

68. Garde R, Dea A, Herwig MF, Ali A, Pincus D. Feedback control of the heat shock response by spatiotemporal regulation of Hsp70. Journal of Cell Biology. 2024;223. doi:10.1083/jcb.202401082

69. Kainth AS, Chowdhary S, Pincus D, Gross DS. Primordial super-enhancers: heat shock-induced chromatin organization in yeast. Trends Cell Biol. 2021;31: 801–813. doi:10.1016/j.tcb.2021.04.004

70. Chowdhary S, Kainth AS, Paracha S, Gross DS, Pincus D. Inducible transcriptional condensates drive 3D genome reorganization in the heat shock response. Mol Cell. 2022;82: 4386–4399.e7. doi:10.1016/j.molcel.2022.10.013

71. Knier AS, Davis EE, Buchholz HE, Dorweiler JE, Flannagan LE, Manogaran AL. The yeast molecular chaperone, Hsp104, influences transthyretin aggregate formation. Front Mol Neurosci. 2022;15. doi:10.3389/fnmol.2022.1050472

72. Winkler J, Tyedmers J, Bukau B, Mogk A. Chaperone networks in protein disaggregation and prion propagation. J Struct Biol. 2012;179: 152–160. doi:10.1016/j.jsb.2012.05.002

73. Rutledge BS, Kim YJ, McDonald DW, Jurado-Coronel JC, Prado MAM, Johnson JL, et al. Stress-inducible phosphoprotein 1 (Sti1/Stip1/Hop) sequesters misfolded proteins during stress. FEBS J. 2024. doi:10.1111/febs.17389

74. Lum R, Tkach JM, Vierling E, Glover JR. Evidence for an unfolding/threading mechanism for protein disaggregation by Saccharomyces cerevisiae Hsp104. J Biol Chem. 2004;279: 29139–46. doi:10.1074/jbc.M403777200

75. Tariq A, Lin J, Noll MM, Torrente MP, Mack KL, Murillo OH, et al. Potentiating Hsp104 activity via phosphomimetic mutations in the middle domain. FEMS Yeast Res. 2018;18: foy042. doi:10.1093/femsyr/foy042

76. Shorter J. The Mammalian Disaggregase Machinery: Hsp110 Synergizes with Hsp70 and Hsp40 to Catalyze Protein Disaggregation and Reactivation in a Cell-Free System. PLoS One. 2011;6: e26319. doi:10.1371/journal.pone.0026319

77. Higuchi-Sanabria R, Pernice WMA, Vevea JD, Alessi Wolken DM, Boldogh IR, Pon LA. Role of asymmetric cell division in lifespan control in Saccharomyces cerevisiae. FEMS Yeast Res. 2014;14: 1133–46. doi:10.1111/1567-1364.12216

78. Kaganovich D, Kopito R, Frydman J. Misfolded proteins partition between two distinct quality control compartments. Nature. 2008;454: 1088–1095. doi:10.1038/nature07195

79. Samant RS, Livingston CM, Sontag EM, Frydman J. Distinct proteostasis circuits cooperate in nuclear and cytoplasmic protein quality control. Nature. 2018;563: 407–411. doi:10.1038/s41586-018-0678-x

80. Malinovska L, Kroschwald S, Munder MC, Richter D, Alberti S. Molecular chaperones and stress-inducible protein-sorting factors coordinate the spatiotemporal distribution of protein aggregates. Mol Biol Cell. 2012;23: 3041–3056. doi:10.1091/mbc.e12-03-0194

81. Mosser DD, Caron AW, Bourget L, Meriin AB, Sherman MY, Morimoto RI, et al. The chaperone function of hsp70 is required for protection against stress-induced apoptosis. Mol Cell Biol. 2000;20: 7146–59. doi:10.1128/MCB.20.19.7146-7159.2000

82. Ben-Zvi A, Miller EA, Morimoto RI. Collapse of proteostasis represents an early molecular event in *Caenorhabditis elegans* aging. Proceedings of the National Academy of Sciences. 2009;106: 14914–14919. doi:10.1073/pnas.0902882106

83. Walther DM, Kasturi P, Zheng M, Pinkert S, Vecchi G, Ciryam P, et al. Widespread Proteome Remodeling and Aggregation in Aging C. elegans. Cell. 2015;161: 919–932. doi:10.1016/j.cell.2015.03.032

84. Moreno DF, Jenkins K, Morlot S, Charvin G, Csikasz-Nagy A, Aldea M. Proteostasis collapse, a hallmark of aging, hinders the chaperone-Start network and arrests cells in G1. Elife. 2019;8. doi:10.7554/eLife.48240

85. David DC, Ollikainen N, Trinidad JC, Cary MP, Burlingame AL, Kenyon C. Widespread Protein Aggregation as an Inherent Part of Aging in C. elegans. PLoS Biol. 2010;8: e1000450. doi:10.1371/journal.pbio.1000450

86. Reis-Rodrigues P, Czerwieniec G, Peters TW, Evani US, Alavez S, Gaman EA, et al. Proteomic analysis of age-dependent changes in protein solubility identifies genes that modulate lifespan. Aging Cell. 2012;11: 120–7. doi:10.1111/j.1474-9726.2011.00765.x

87. Feder JH, Rossi JM, Solomon J, Solomon N, Lindquist S. The consequences of expressing hsp70 in Drosophila cells at normal temperatures. Genes Dev. 1992;6: 1402–1413.

88. Huh W-K, Falvo J V., Gerke LC, Carroll AS, Howson RW, Weissman JS, et al. Global analysis of protein localization in budding yeast. Nature. 2003;425: 686–691. doi:10.1038/nature02026

89. Gietz RD, Schiestl RH. High-efficiency yeast transformation using the LiAc/SS carrier DNA/PEG method. Nat Protoc. 2007;2: 31–34. doi:10.1038/nprot.2007.13

90. Li B, Dewey CN. RSEM: accurate transcript quantification from RNA-Seq data with or without a reference genome. BMC Bioinformatics. 2011;12: 323. doi:10.1186/1471-2105-12-323

91. Dobin A, Gingeras TR. Mapping RNA-seq Reads with STAR. Curr Protoc Bioinformatics. 2015;51: 11.14.1-11.14.19. doi:10.1002/0471250953.bi1114s51

92. Love MI, Huber W, Anders S. Moderated estimation of fold change and dispersion for RNA-seq data with DESeq2. Genome Biol. 2014;15: 550. doi:10.1186/s13059-014-0550-8

93. Raudvere U, Kolberg L, Kuzmin I, Arak T, Adler P, Peterson H, et al. g:Profiler: a web server for functional enrichment analysis and conversions of gene lists (2019 update). Nucleic Acids Res. 2019;47: W191–W198. doi:10.1093/nar/gkz369

94. Schindelin J, Arganda-Carreras I, Frise E, Kaynig V, Longair M, Pietzsch T, et al. Fiji: an open-source platform for biological-image analysis. Nat Methods. 2012;9: 676–682. doi:10.1038/nmeth.2019

95. Garde R, Singh A, Ali A, Pincus D. Transcriptional regulation of Sis1 promotes fitness but not feedback in the heat shock response. Elife. 2023;12. doi:10.7554/eLife.79444

96. Neferkara A, Ali A, Pincus D. Cellquant: a vibecoder’s guide to image analysis. 2026. doi:10.64898/2026.03.09.710634

97. Petropavlovskiy AA, Tauro MG, Lajoie P, Duennwald ML. A Quantitative Imaging-Based Protocol for Yeast Growth and Survival on Agar Plates. STAR Protoc. 2020;1: 100182. 10.1016/j.xpro.2020.100182

98. Pau G, Fuchs F, Sklyar O, Boutros M, Huber W. EBImage-an R package for image processing with applications to cellular phenotypes. BIOINFORMATICS APPLICATIONS NOTE. 2010;26: 979–981. doi:10.1093/bioinformatics/btq046

99. von der Haar T. Optimized Protein Extraction for Quantitative Proteomics of Yeasts. PLoS One. 2007;2: e1078. doi:10.1371/journal.pone.0001078

