## Supplementary material for "Mitochondrial respiration modulates Hsf1 activation and the heat shock response": Fig. S1

### SUPPLEMENTARY FIGURES

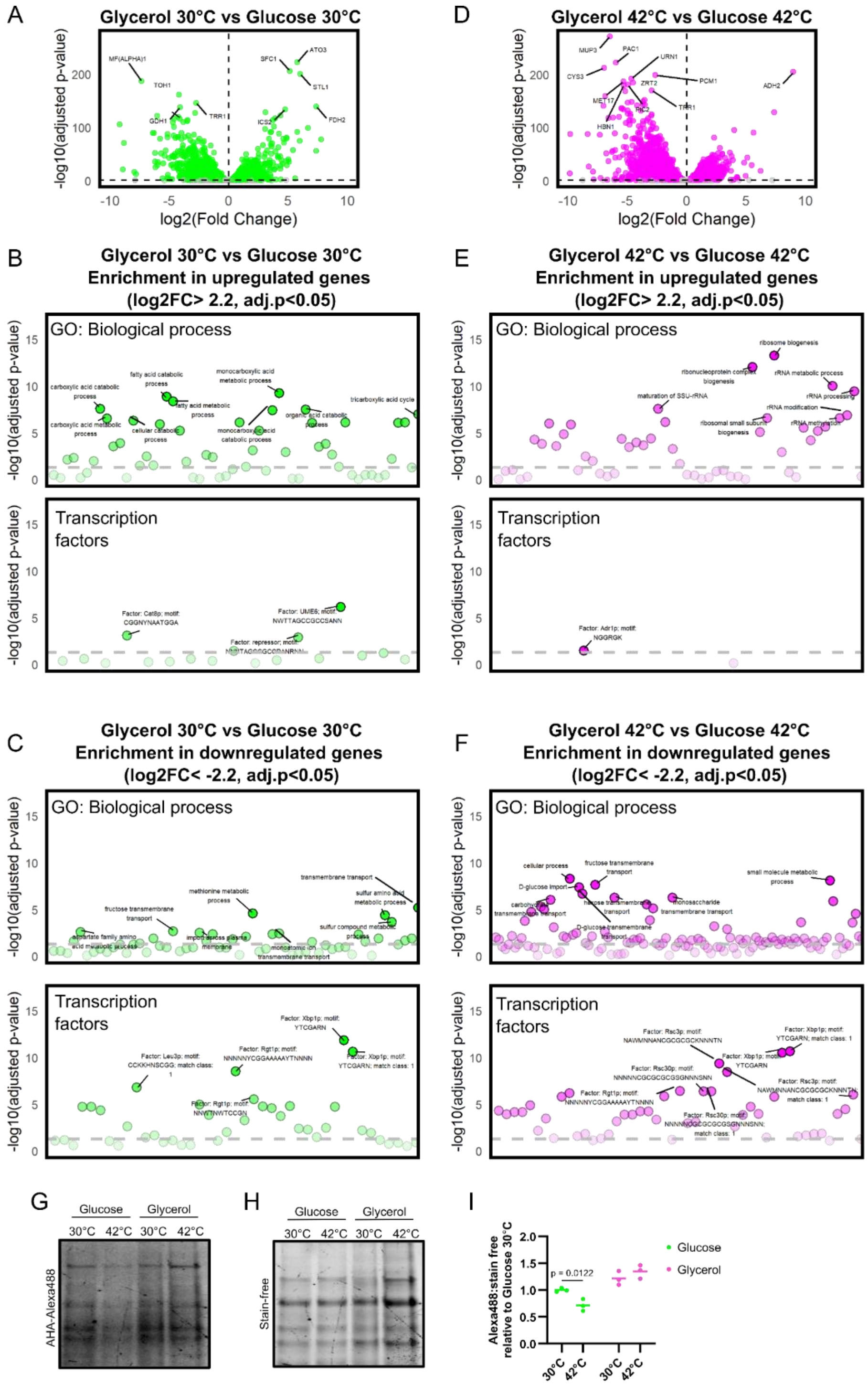

**Figure S1: Respiring cells upregulate ribosome-associated transcripts in response to heat shock.** A. Volcano plot showing upregulated and downregulated genes in cells grown in glycerol at 30°C relative to cells grown in glucose at 30°C. B. Enriched GO terms (Biological processes) and transcription factors associated with the set of upregulated genes (log2FC>2.2) in cells grown in glycerol at 30°C relative to cells grown in glucose at 30°C. C. Enriched GO terms (Biological processes) and transcription factors associated with the set of downregulated genes (log2FC<-2.2) in cells grown in glycerol at 30°C relative to cells grown in glucose at 30°C. D. Volcano plot showing upregulated and downregulated genes in cells grown in glycerol and heat shocked at 42°C for 30 mins relative to cells grown in glucose and heat shocked at 42°C for 30 mins. E. Enriched GO terms (Biological processes) and transcription factors associate with the set of upregulated genes (log2FC>2.2) in cells grown in glycerol and heat shocked at 42°C for 30 mins relative to cells grown in glucose and heat shocked at 42°C for 30 mins. F. Enriched GO terms (Biological processes) and transcription factors associated with the set of downregulated genes (lof2FC<-2.2) in cells grown in glycerol and heat shocked at 42°C for 30 mins relative to cells grown in glucose and heat shocked at 42°C for 30 mins. G. SPAAC pulse-labelled de novo synthesized proteins and H. total protein on stain-free gel of lysates extracted from cells grown in glucose and glycerol before and after a heat shock at 42°C for 1 hour.
